# An effective, safe and cost-effective cell-based chimeric vaccine against SARS-CoV2

**DOI:** 10.1101/2020.08.19.258244

**Authors:** Zhenguo Cheng, Danhua Zhang, XiaoWen Liu, Jinxin Miao, Jianyao Wang, Haoran Guo, Wenli Yan, Zhe Zhang, Na Zhang, Jingjing Wang, Shuangshuang Lu, Zhongxian Zhang, Wei Liu, Hong Liu, Yi Zhang, Lirong Zhang, Jianzeng Dong, Nicholas R Lemoine, Yaohe Wang

**Affiliations:** Sino-British Research Centre for Molecular Oncology, National Centre for International Research in Cell and Gene Therapy, School of Basic Medical Sciences, Academy of Medical Sciences, Zhengzhou University, Zhengzhou 450052, China; State Key Laboratory of Esophageal Cancer Prevention & Treatment, Zhengzhou University 450052, Zhengzhou, China; The First Affiliated Hospital of Zhengzhou University, Zhengzhou University, Zhengzhou, 450052, Henan; The Affiliated Children’s Hospital of Zhengzhou University, Zhengzhou University, Zhengzhou, 450018, China; Department of Pharmacology, School of Basic Medical Sciences, Zhengzhou University, Zhengzhou, Henan 45001, China; Centre for Biomarkers & Biotherapeutics, Barts Cancer Institute, Queen Mary University of London, London, EC1M 6BQ, UK

**Keywords:** SARS-CoV2, chimeric vaccine, spike, nucleocapsid, neutralizing antibody, T cell immunity

## Abstract

More than one hundred vaccines against SARS-CoV-2 have been developed and some of them have entered clinical trials, but the latest results revealed that these vaccines still face great challenges. Here, we developed a novel cell-based gp96-Ig-secreting chimeric vaccine which is composed of two viral antigens, the RBD of spike protein, and a truncated nucleocapsid protein that could induce epitope-specific cytotoxic T lymphocytes but low antibody response. Syrian hamsters immunized with the cell-based vaccine produced high level of SARS-CoV-2 specific NAbs and specific T cell immunity which could eliminate RBD-truncated N-expressing cells, without the induction of antibody against N protein and other observed toxicity. This study provides a proof of concept for clinical testing of this safe, effective and cost-effective vaccine against SARS-CoV2 infection.

---

SARS-CoV2 is a novel Sarbecovirus subgenus of coronavirus which has caused a world epidemic of the coronavirus-induced disease 19 (COVID-19) (*1*). As of August 6, more than 18 million people have been diagnosed, with 701,544 confirmed deaths (*2*). Although several effective neutralizing antibodies have been identified for clinical interventions against SARS-CoV-2 infection (*3*), an efficient vaccine againsts COVID-19 is still not available. Currently, there are 25 candidate vaccines in clinical evaluation or in clinical trial, and another 139 candidate vaccines are in preclinical evaluation. Even though the latest Phase I and Phase II clinical trials of adenovirus vectored vaccines have shown encouraging results (*4, 5*), vaccines only targeting Spike protein might not be the best possible designs, and improved versions are needed.

The surface glycoprotein spike (S) of SARS-CoV mediates the attachment and entry of virion into the host cell by binding to its receptor (*6*). Similar to SARS-CoV, SARS-CoV-2 S protein also adopts human angiotensin-converting enzyme 2 (ACE2) as primary receptor (*1*), and possesses a higher affinity (*7*). An antibody against the spike that prevents the virus from getting into cells will theoretically stop the virus from causing disease, therefore S protein is an ideal target for vaccine and antiviral therapy (*8*). However, recent data showed that the serum from COVID-19 patients contained high titer S1-specific antibodies, while antibodies against receptor binding domain (RBD) were low (*9*). Moreover, a subset of patients infected with SARS-CoV-2 might not develop long-lasting antibodies to the virus, and S-antibodies were reported to rapidly decline for people infected with seasonal coronaviruses and who recovered from COVID-19, especially those with mild symptoms or asymptomatic infection (*10*). Furthermore, although there are inconclusive data to prove the presence of antibody-dependent enhancement (ADE) for SARS-COV-2 infection, it is important to consider ADE in the context of efforts to develop countermeasures against the SARS-CoV-2 (*11*), as the data from previous SARS-CoV research strongly suggest that some antibodies that bind viral S protein ADE can facilitate uptake by human macrophages and B cells via their Fcγ receptors (FcγRs) (*12*).

The S protein on the surface of coronavirus is composed of S1 and S2 subunits (Fig. 1A). Binding of the RBD domain in S1 to ACE2 triggers a conformational change of the homotrimer, in which the resulting S2 subunit further forms a six-helix bundle and completes the fusion of virus to host cell membrane (*13*). Our analysis showed that the homology of the S1 subunit between SARS-CoV and SARS-CoV-2 is lower than for the S2 subunit (67.1% vs 90%), and residues located at RBD varied greatly (Fig. S1A&B), indicating that vaccine based on S alone could be challenged to prevent infection with SARS-CoV or some SARS-CoV2 mutants. Given the importance of the homotrimer structure of S protein for the induction of neutralising antibody, we searched for potential immunogenic epitopes of S protein based on its 3D structure using DiscoTope. As shown in Fig 1B, the predicted 3D immunogenic epitopes were fewer compared to those predicted based on linear sequence (Fig. 1C). Moreover, these predicted immunogenic 3D epitopes were located in the loop region of RBD which is also the main interface between RBD and ACE2 (Fig. 1D). Of note, it was also found that the RBD of SARS-CoV-2 had weaker immunogenicity than that from SARS-CoV (Fig. S1C&D), which may be a reason why SARS-CoV-2 is so widely spread. Abundant studies demonstrate that different codon usage bias plays an important role in protein translation (*14*). We found that the protein expression of the S gene using original sequence was almost undetectable in human 293T cells, while the optimised codon significantly increased the S protein expression (Fig. 1E&F), indicating a codon-optimized S gene is necessary for SARS-COV-2 vaccine. Recently, it has been demonstrated that the receptor-binding domain of SARS-CoV-2 elicits a potent neutralizing response without ADE (*15*), suggesting RBD as a potential candidate for the development of new vaccine.

**Figure 1.**
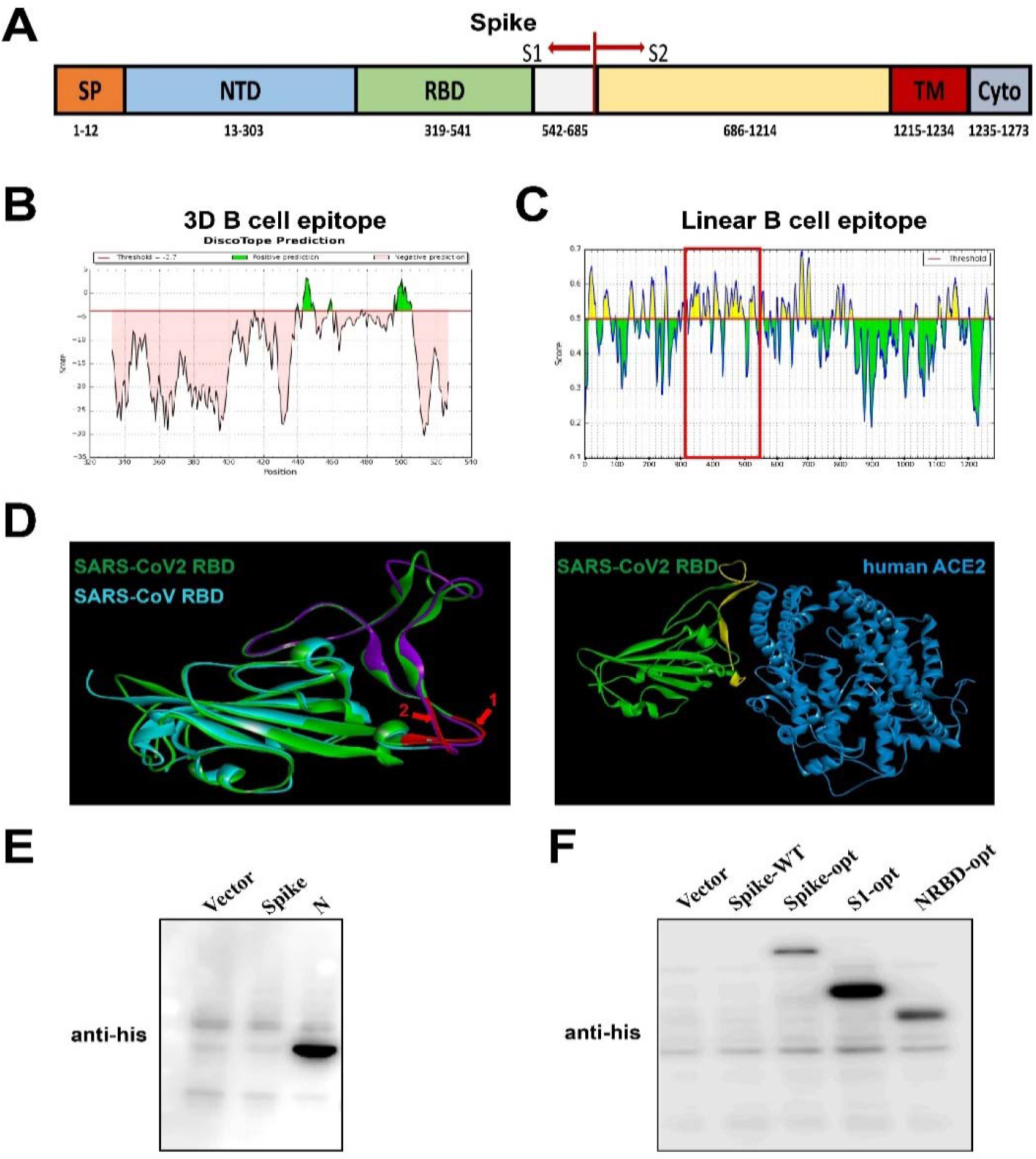
The RBD domain of Spike is crucial for the SARS-CoV2 Vaccine. (A) The functional domain of SARS-CoV-2 spike protein. (B) Potential B cell antigen of RBD domain from SARS-CoV2 is predicted by Discotope software based on their 3D structure. (C) Potential linear B cell epitopes of SARS-CoV-2 full S protein are analysed with the IEDB database. (D) The location of potential antigens in RBD domain (SARS-CoV-2:red, SARS-CoV: Purple) and interaction model between RBD and ACE2 receptor (interface is marked yellow) are marked with Discovery Studio. (E) The expression of Spike and nucleocapsid with wild-type sequence in 293T cells are detected by Western Blot assay. (F) The expression of Spike and its derivatives with codon optimization (opt).

Extensive research into animal and human SARS-CoVs have demonstrated that both neutralising antibody and T-cell responses are important in eliminating the virus and controlling disease development of SARS-CoV-2 infection (*16*). The latest clinical data has shown that there were abundant SARS-CoV2-specific T cells in PBMC from convalescent patients (*17*), indicating that enhancing specific antiviral T cell responses will be helpful for preventing SARS-CoV-2 infection. Nucleocapsid (N) protein is a potential candidate for induction of T cell immunity. N protein, the most abundant and highly conserved protein of coronavirus, is involved in virus assembly by packaging the viral genome into a ribonucleocapsid (RNP) (*18*). Antibodies against N protein in patients’ serum have been used as a diagnosis marker for SARS-CoV2(*19, 20*). Previous findings about SARS had shown that vaccination with N protein could inhibit virus infection by inducing specific antiviral antibodies and T cell immune responses (*16, 21*). However, a study showed that the vaccine based on full N protein significantly induced inflammatory cell infiltration and pathogenic anti-N antibody, which exacerbates lung injury in mice (*22, 23*). Consistent with this finding, a recent serological study uncovered that a higher titre of anti-N IgG and IgM was correlated with worse clinical readout of COVID-19 patients (*24*). In addition, inactivated virus vaccine could induce strong pathological N antibodies, which will also make it difficult to distinguish the truly infected and vaccine-immunised populations (*21, 25*). Therefore, identification of the peptides of N protein with low antibody induction and high cytotoxic T lymphocytes stimulation will be a game-changer to design a safer and effective vaccine to overcome the bottleneck of N protein-based vaccine. To this end, a homology analysis and potential epitope prediction were performed. As shown in Fig S. 1E, N protein shares 91.2% amino acid identity between SARS-CoV-2 and SARS-CoV, which supports the results that convalescent sera from SARS-CoV have high cross-reactivity with emerging SARS-CoV-2 (*26*). Epitope analysis using the IEDB database demonstrated that there were two regions in the N protein that had fewer B-cell epitopes but abundant MHC I-binding T-cell epitopes (Fig. 2A &B). Moreover, previous serological research into SARS had demonstrated that the antibodies against peptides 212-341aa were less prevalent in SARS patients (Fig. 2C) (*27*). Accordingly, we constructed a chimeric vaccine which was composed of NAb-inducing Spike RBD and a truncated N protein only containing T cell-associated peptide (Ntap) (Fig. 2D). To validate this hypothesis, the plasmids containing the chimeric vaccine or its derivatives were constructed and transfected into 293T cells. As predicted, convalescent sera from six COVID-19 patients had highly reactive antibodies against full N protein, while antibodies that recognized S protein and the Ntap region were poor (Fig. S2). Strikingly, the truncated N protein was not recognised by the commercial polyclonal antibodies and convalescent sera from COVID-19 patients (Fig. 2E). Most importantly, two epitope peptides (LLLDRLNQL and GMSRIGMEV) from our identified truncated N proteins had a high binding affinity to the HLA-A*0201 molecule (Fig. 3A) while peptide RLNQESKM from SARS-COV-2, which had a mutation compared with SARS-CoV (RLNQESKV), was less able to the HLA-A*0201 molecule. Encouragingly, the latest studies found that the memory T cells from SARS-CoV-recovered patients exhibited long-lasting T cell immunity and robust cross-reactivity to SARS-CoV-2 N protein (*28*), which also supports the superiority of our strategy.

**Figure 2.**
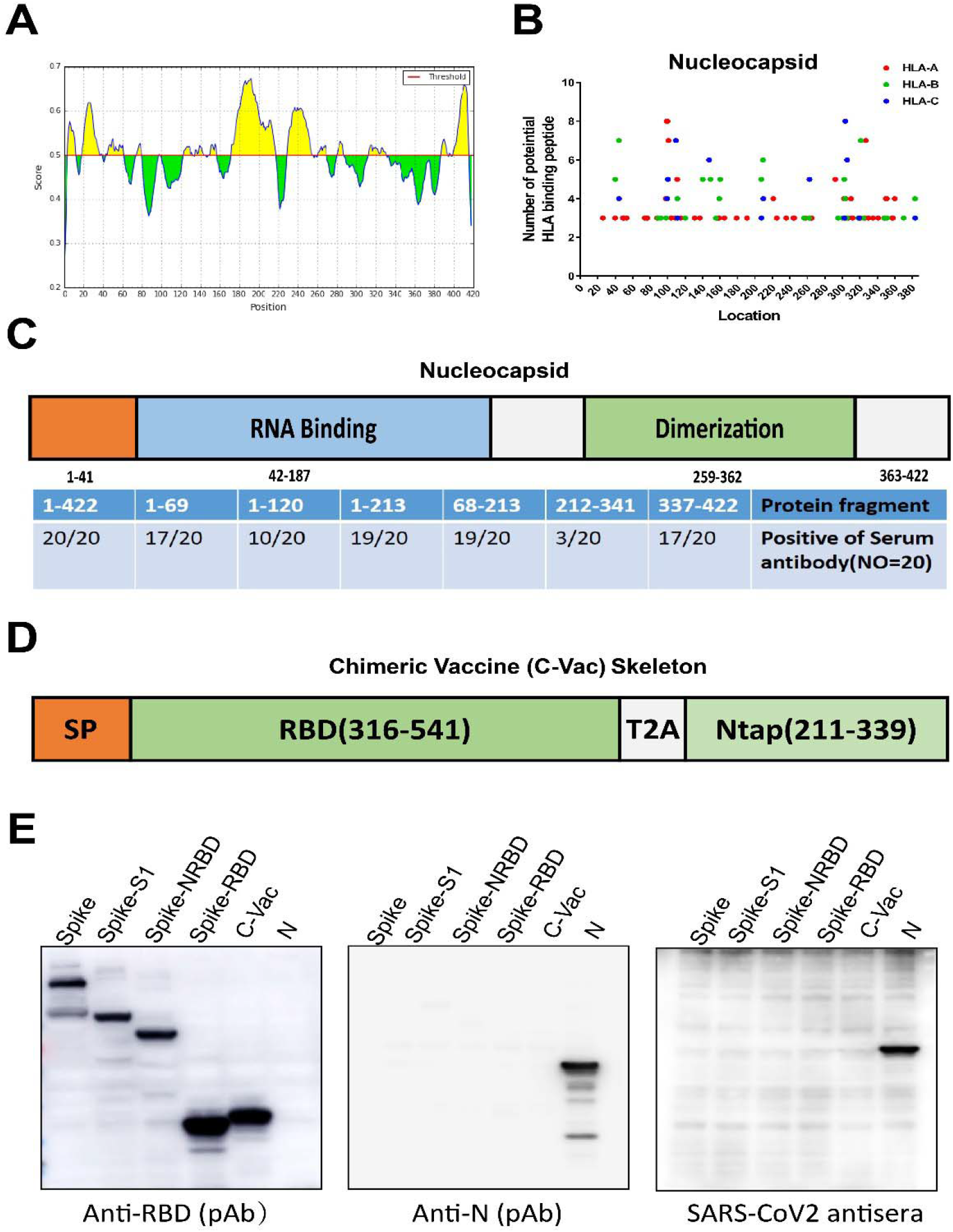
Construction of chimeric vaccine for SARS-CoV-2. (A) Potential B-cell epitopes of N protein is predicted by IEDB database. (B) Potential MHCI-binding peptides of N. (C) Functional domain of SARS-CoV N protein (Upper) and its antibody epitope map reported in previous study. (D) The skeleton of Chimeric Vaccine for SARS-CoV-2, RBD: spike RBD domain (306-541 aa), Ntap: T-cell-associated peptide of N (211-339 aa). (E) Characterization of SARS-CoV-2-derived protein and C-Vac antigen by SARS-CoV-2 antisera and commercial antibodies against SARS-CoV2 spike RBD or Nucleocapsid.

**Figure 3.**
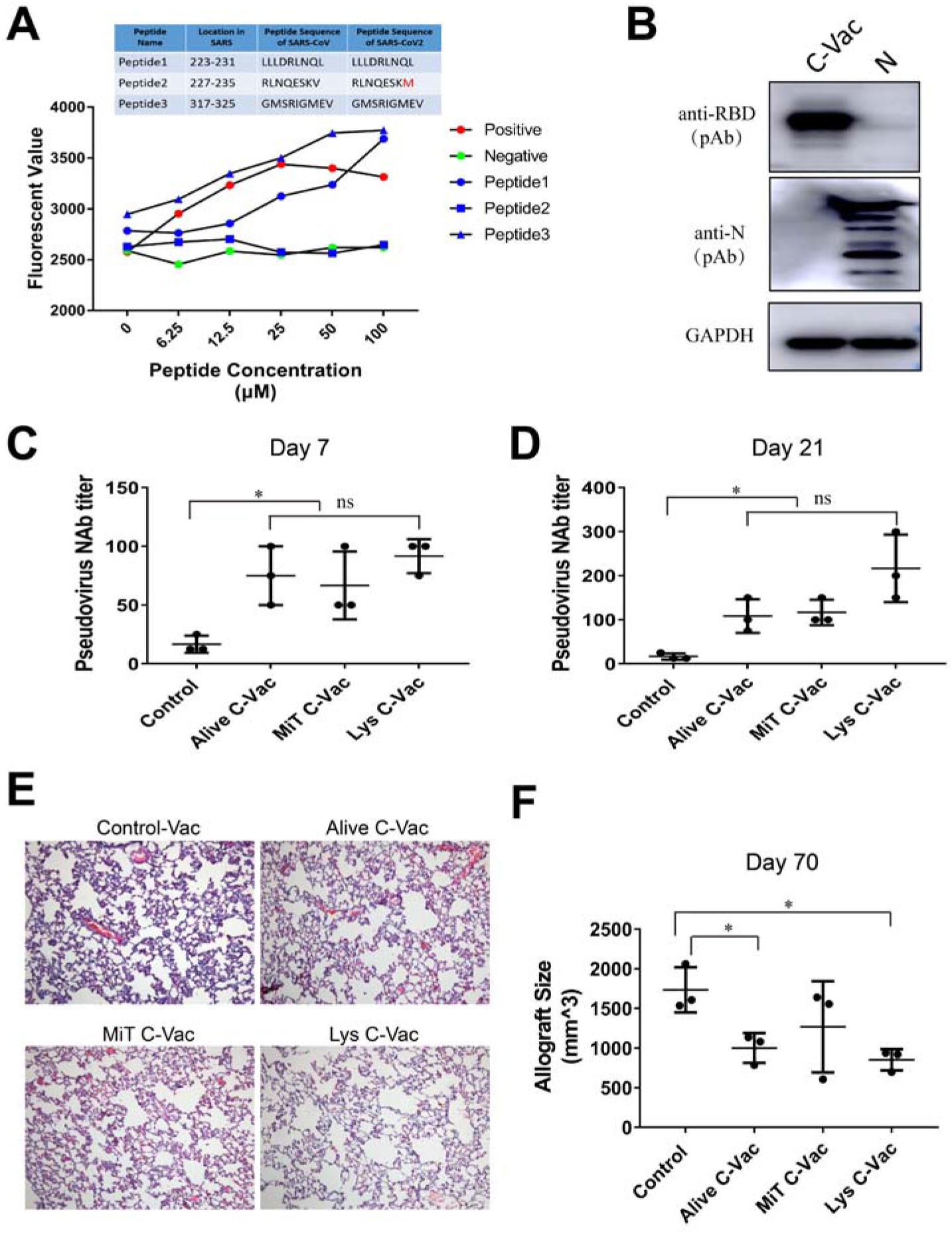
The immunogenicity, efficacy and safety of C-Vac for SARS-CoV-2 infection. (A) Flow cytometry-based detection of MHC(HLA-A2)-peptide complex binding affinity in T2 cells. (B) The expression of antigens in 293T-based C-Vac is confirmed by Western Blot. N, 293T cells transfected with the plasmid expressing a full length N gene. (C&D) Pseudovirus neutralization titers of hamster serum at day 7 after the first immunization and day 21 (boosted at day 14) after vaccination with 293T-based C-Vac, MiT C-Vac: Mitomycin C-treated C-Vac, Lys C-Vac: Lysed C-Vac. (E) Histological characteristics of hamster lung at day 7 after the first vaccination. Original magnification 200× (F) Allograft volume of transformed fibroblasts expressing RBD-truncated N protein in the Syrian hamsters immunized with different regime. 5×10^6^ BHK21 cells expressing C-Vac antigen (RBD-Ntap) were subcutaneously injected into immunized hamsters for challenge at day 45 after boost, and the volume of allografts were measured at 14 days after inoculation of the BHK21 cells expressing RBD-Ntap into the immunized hamsters.

Having identified more effective and safer promising antigens, next we chose an efficient and largely scalable production approach to deliver our optimised chimeric RBD-Ntap with low cost and rapid GMP production. Cell-based gp96-Ig-secreting vaccines have been reported as a potent modality to induce both systemic and mucosal immunity for cancer and infectious diseases (*29*). In order to enhance the efficiency of this vaccine, we constructed a gp96-Ig-secreting chimeric cell vaccine (C-Vac) using the human 293T cell line stably expressing RBD-Ntap (Fig.3B and Fig. S1F), given that 293T cells express adenovirus early proteins and SV40 (Simian Virus 40) large antigen protein, which are potential danger signals resulting in strong immunity. Since the Syrian hamster has been demonstrated to be permissive to SARS-CoV-2 infection (*30*), we also validated that ACE2 receptor from both human and hamster could support pseudovirus infection (data not shown), indicating that the hamster is a suitable animal model to evaluate the efficacy and safety of SARS-CoV2 vaccine. To assess the efficacy and safety of our C-Vac, Syrian hamsters were immunized with a series of 293T-based vaccines (untreated live cells, mitomycin C-treated cells and freeze-thaw cell lysis), then the NAbs in the vaccinated hamsters were evaluated at different times points by a pseudovirus infection assay using human ACE2-overexpressing Syrian hamster BHK21 cells as indicator cells. All three vaccinated groups could effectively induce NAbs at day 7 with single immunisation (Fig 3C) and at day 21 with double immunization (boost was performed on day 14) (Fig 3D), and had no lung injury observed in any of these animals (Fig 3E).

Interestingly, our results showed that the freeze-thaw-treated 293T cell vaccine appear somewhat better than live or mitomycin C-treated 293T cell vaccines although there was no significant difference (p>0.05). In order to prove the efficacy of C-Vac further, Syrian hamster-derived BHK-21 fibroblasts expressing RBD-Ntap (C-Vac antigen) were injected subcutaneously into the Syrian hamsters immunised with different vaccination regimes as indicated. Strikingly, the hamsters immunized with live and lyzed C-Vac significantly reduced the allograft volume (Fig. 3F), indicating that our C-Vac could eliminate RBD-Ntap expressing cells due to viral antigen-specific T cell immunity. All these data imply that our C-Vac lysed cell vaccine is a rapid, convenient and cost-effective vaccine for SARS-CoV-2, as amplification of 293T cells is much easier and less costly compared to other approaches, such as inactivated virus vaccine, viral vector vaccine and RNA vaccines.

Now it is urgent to develop an effective and safer vaccine to tackle the spread of COVID-19 pandemic worldwide. Although a large number of vaccines against SARS-CoV-2 are undergoing development in preclinical studies and clinical trials, most vaccines just target the spike protein. In this study, we developed a completely new and promising cell-based chimeric vaccine (C-Vac) to target both the RBD of S protein and Nucleocapsid (N). This vaccine could effectively induce NAbs and specific CTLs against viral proteins, but has low potential risk of inducing ADE and anti-N protein antibody-mediated immunotoxicity. Our C-Vac offers the potential for recipients to develop long-lasting immunity to the virus, and will provide a poteintial second defence against SARS-CoV-2 mutants through targeting the most conserved N protein. Due to the lack of P3 laboratory for working with live SARS-COV-2, the C-Vac vaccine developed in this study is currently not able to be tested by live SARS-CoV2 challenge in suitable animal models. However, this study provides a solid proof of concept for clinical testing of this safer, effective and cost-effective vaccine against SARS-CoV-2 infection.

## Materials and Methods

### Plasmids

The coding sequence of Spike, Nucleocapsid derived from SARS-COV2 (MN908947.3), and human IgG Fc fragment (BC065820.1) was synthesized by Sangon Biotech, codon optimized SARS-CoV2 spike was bought from Sino Biological. Sequence of human ACE2 and secreting gp96 (without “KDEL” motif of C terminal) were amplified from Suit2 and 293T cell respectively. Plasmids including 3.1-his-S, 3.1-his-N, 3.1-his-optS, 3.1-his-optS1, 3.1-his-optNRBD (from N terminal to RBD domain), 3.1-optS-d18 (without the last 18 amino acid residues) and 3.1-gp96-hFc were cloned into pcDNA3.1-his tag (hygro) vector with the following primer. Spike-F: ACTTAATTAAGCCACCATGTTTGTTTTTCTTGTTTTATTGCCAC, Spike-R: TAACCGGTTGTGTAATGTAATTTGACTCCTTTGAGC; Nucleocapsid-F: ACGGATCCGCCACCATGTCTGATAATGGACCCCAAAATCAG, Nucleocapsid-R: ACGTACCTGCAGGACCGGTGCTAGCATTAATGATGATGATGATGATGACCG; Spike-optF (codon optimized): ACTTAATTAAGCCACCATGTTTGTGTTCCTGGTGCTGCT, Spike-opt-R: TAACCGGTGGTGTAGTGCAGTTTCACTCCTTTCA; Spike-optS1-R: TAACCGGTGCTGTTGGTCTGGGTCTGGTAGG; Spike-optNRBD-R: TAACCGGTTCCATTGAAGTTGAAGTTCACACACTT; Spike-optSd18: ATACCGGTTTACTTACAACAGGAGCCACAGGAAC; gp96-F: TACTCGAGGCTGACGATGAAGTTGATGTGGA, gp96-R: TAGCTAGCTTCAGCTGTAGATTCCTTTGCTGTTTC; hFc-F: TAGCTAGCGAGCCCAAATCTTGTGACAAAACTCA, hFc-R: ACGGTACCACGCGTTTATTTACCCGGAGACAGGGAGAGGCTC. The chimeric vaccine with codon optimized RBD domain (containing signaling peptide), T2A cleavage peptide and truncated N was synthesized (Sangon Biotech) and inserted into modified pLenti6-puromycin vector. Lentivirus plasmids containing human ACE2 or luciferase reporter gene were generated with the following primers, hACE2-F: ACTTAATTAAGCCACCATGTCAAGCTCTTCCTGGCTCCT, hACE2-R: ATACCGGTAAAGGAGGTCTGAACATCATCAGTG; luc-F: ATACCGGTGCCACCATGGAAGACGCCAAAAACATAAAG, luc-R: ATGTCGACTTACACGGCGATCTTTCCGC. Lentivirus packing helper plasmids psPAX (#12260) and pMD2.G (#12259) were provided by addgene.

### Cells

Human kidney cell line HEK-293T and hamster kidney cell line BHK21 were purchased from Cell Bank of Type Culture Collection Committee of Chinese Academy of Sciences. Cells were cultured in Dulbecco’s Modified Eagle Medium (DMEM high glucose, Gibco) with 10% FBS (Biological Industries) and 2 mM GlutaMax™ (Gibco). T2 cell (ATCC) that does not express HLA DR and are Class II major histocompatibility (MHC) antigen negative was maintained in Iscove’s Modified Dulbecco’s Medium (IMDM, Invitrogen) supplemented with 20% FBS. 293T-gp96-hFc overexpressing cell line was constructed by transfected 3.1-gp96-hFc plasmid with Higene (Applygen, China) and selected with 200μg/ml hygromycin B (Invitrogen) according to the manufacturer’s instruction. To generate BHK21-hACE2 or C-Vac overexpressing cells, 5×104 cells (500 μL) BHK21 or 293T-gp96-hFc cells were seeded into 24-well plates, then 1 ml supernatant containing proper lentivirus and 5 μg/ml polybrene (Sigma) was added. 12 h later, medium was refreshed, and 5 μg/ml puromycin (Selleck) was added to screen the stable cell lines 72 h after infection.

### Protein isolation and western blot assay

Cells were washed twice with ice-cold phosphate-buffered saline and lysed on ice for 10 min with RIPA lysis buffer (50 mM Tris • HCl, pH 7.4, 0.1% SDS, 150 mM NaCl, 1 mM EDTA, 1 mM EGTA, 10 mM NaF) containing 1% protease inhibitor cocktail solution (Roche). Cell lysates were clarified by centrifugation at 13000 rpm for 30 min at 4°C, then 25 ug total protein was separated by 10% SDS-PAGE and transferred to PVDF membranes (Millipore). Membranes were blocked with 5% non-fat milk and probed with the indicated primary antibodies following the manufacturer’s instruction. After incubation with horseradish peroxidase-conjugated secondary antibodies (1:5000, ZSBIO) for 1 h at room temperature, membranes were washed three times with TBST and detected by the enhance chemiluminescence (ECL) system (Thermo pierce, USA). Antibodies used in this study were listed as the following, rabbit anti-RBD PAb (polyclonal antibody) (Sino Biological, #40592-T62), rabbit anti-Nucleocapsid PAb (Sino Biological, #40588-T62), mouse anti-GAPDH mAb (monoclonal antibody) (ProteinTech, 60004-1-Ig), mouse anti-his tag mAb (Abmart, M20001S), HRP Goat Anti-Rabbit IgG (ZSBIO, ZB-5301), HRP Goat anti-mouse IgG (ZSBIO, ZB-5305), HRP Goat anti-human IgG (ZSBIO, ZB-2304).

### Cell-based peptide MHC affinity assay

T2 cell which has no expression of HLA DR and Class II major histocompatibility (MHC) antigen was used to evaluate the affinity between peptide and HLA A*0201. Briefly, 100 μL 1×105 T2 cells were seeded into round bottom 96-well plate with FBS free IMEM medium, and charged with multiply dilution peptides from the maximal concentration of 100 μM for 4 h. Then cells were washed twice with cold PBS and stained with mouse anti-human HLA-A2-FITC antibody (Abcam) for 30 min on ice. Subsequently, the fluorescence intensity of cells were analyzed using a BD FACSAria (BD Biosciences Immunocytometry Systems) after washed twice with cold PBS. All peptides including three potential HLA A*0201 binding peptides from N (LALLLLDRLNQL, RLNQLESKM, GMSRIGMEV), one positive peptide (KIFGSLAFL) derived from HER2 derived positive peptide and a negative control peptides (SAPDTRPAP) from MUC1 were synthesized by GL Biochem.

### Serum samples from recovered COVID-19 patients

6 serum samples from recovered COVID-19 patients were collected in the First Affiliated Hospital of Zhengzhou University with the patients’ written consent and was approved by the Ethics Committee of Zhengzhou University.Samples were heat-inactivated at 56°C and stored in aliquots at −80°C.

### Animals

Female,12-week-old Syrian hamster were purchased from Vitalriver (Beijing), and randomly divided into 4 groups (n=9). Group1-4 were vaccinated with 1×107 293T-gp96-hFc control cells, viable 293T-C-Vac cells, 293T-C-Vac cells treated with 5μg/ml mitomycin (MCE) for 4h and freeze-thaw treated 293T-C-Vac cells lysates, respectively. Three hamsters in each groups were sacrificed at the day 7 post-immunization and animal serum samples were collected and store at −80◻, while the lung tissues were fixed in 10% neutral-buffered formalin and embedded in paraffin. Remaining animals were boosted at day 14 with the same regime as prime, then serum was harvested at day 21 (7 days after boost). Finally, hamster were challenged with hamster BHK21 cells expressing RBD-Ntap (C-Vac antigen) and the volume of allografts were measured to evaluate vaccine-specific cytotoxic T cells. All animals used in this study were maintained under specific pathogen-free conditions in Laboratory Animal Center of Zhengzhou University, and treatments was in accordance with the NIH Animal Care and Use Committee regulations.

### Hematoxylin-Eosin staining and Immunocytochemistry

Hamster lung tissues were sectioned at 6 μm, and resulting slides were stained with hematoxylin and eosin. Briefly, slides were firstly cleared the paraffin using xylene for three times, then hydrated by incubating in 100%, 100%, 95%, 90% and 80% ethanol and water for 5min respectively. After samples were stained with hematoxylin solution for 5 min and eosin Y solution for 2 min, transferring slides into 80%, 90%, 95%, 100%, 100% ethanol for dehydration and in xylene three times for transparency. For Immunocytochemistry, fixed 293T-gp96-hFc or control cells were punched with 1‰ TritonX-100, then incubated with HRP Goat anti-human IgG and stained with DAB kit (Maxim Biotechnologies) following to the instructions. All slides were covered with neutral resin and took photos with microscope (Leica).

### Generation of Lentivirus and SARS-CoV2 pseudovirus

To generate lentivirus expressing hACE2 or C-Vac, 2×106 293T cells were seeded in 10 cm cell culture dish and co-transfected the plasmids mixture including 25μg target plasmid, 8μg psPAX2, 4μg pMD2.G with 40 μL PEI (2mg/ml, Sigma). The medium was refresed after 6 hours, and supernatants containing lentivirus were collected 48 hours later(2). For SARS-CoV2 pseudovirus, 293T cells were co-infected 20μg lenti-Luc, 10μg psPAX and 10μg 3.1-optS-d18 and pseudovirus were harvested. Subsequently, supernatant containing virus was filtered with 0.22μm filters (Millipore) and store at −80◻ in aliquots.

### Pseudovirus neutralization assay

Hamster sera from vaccinated and control groups were heat inactivated for 30 min at 56◻,and serially diluted twofold starting from 1:25. Next, a fix amount of pseudovirus (50μL) was incubated with 50μL diluted sera in 37◻ cell incubator for 1 hour, then 1×104 BHK21 expressing hACE2 were seeded into virus-sera mixture. 72 h after infection, relative luciferase activity of cells were measured with luciferase assay system (Promega) and GloMax Discover detector (Promega). Sera neutralization titers (ID50) were calculated using 50% RLU signaling compared with control cells.

### Bioinformatics analysis

Protein homology analysis between SARS-CoV and SARS-CoV2 were performed by MegAlign software. Potential B cell epitopes based on 3D structure or linear sequence and MHC-I binding peptides were analyzed with IEDB database (http://tools.iedb.org). The RBD domain from SARS-CoV2 (PDB: 6LZG) and SARS-CoV (PDB: 2GHW) were compared with 3D-Match (http://www.softberry.com/berry.phtml) and displayed using Discovery Studio.

### Statistics analysis

All statistical tests were performed by Graphpad Prism 7 using Student’s t-test (unpaired, two-tailed).The differences were considered statistically significant with a p-value < 0.05.

## Acknowledgement

This study was supported by grants from the National Natural Science Foundation of China (81702383, U1704282) and National Key Technologies R & D Program of China (2016YFE0200800).

