## Supplemental figures for "An effective, safe and cost-effective cell-based chimeric vaccine against SARS-CoV2"

Fig. S1

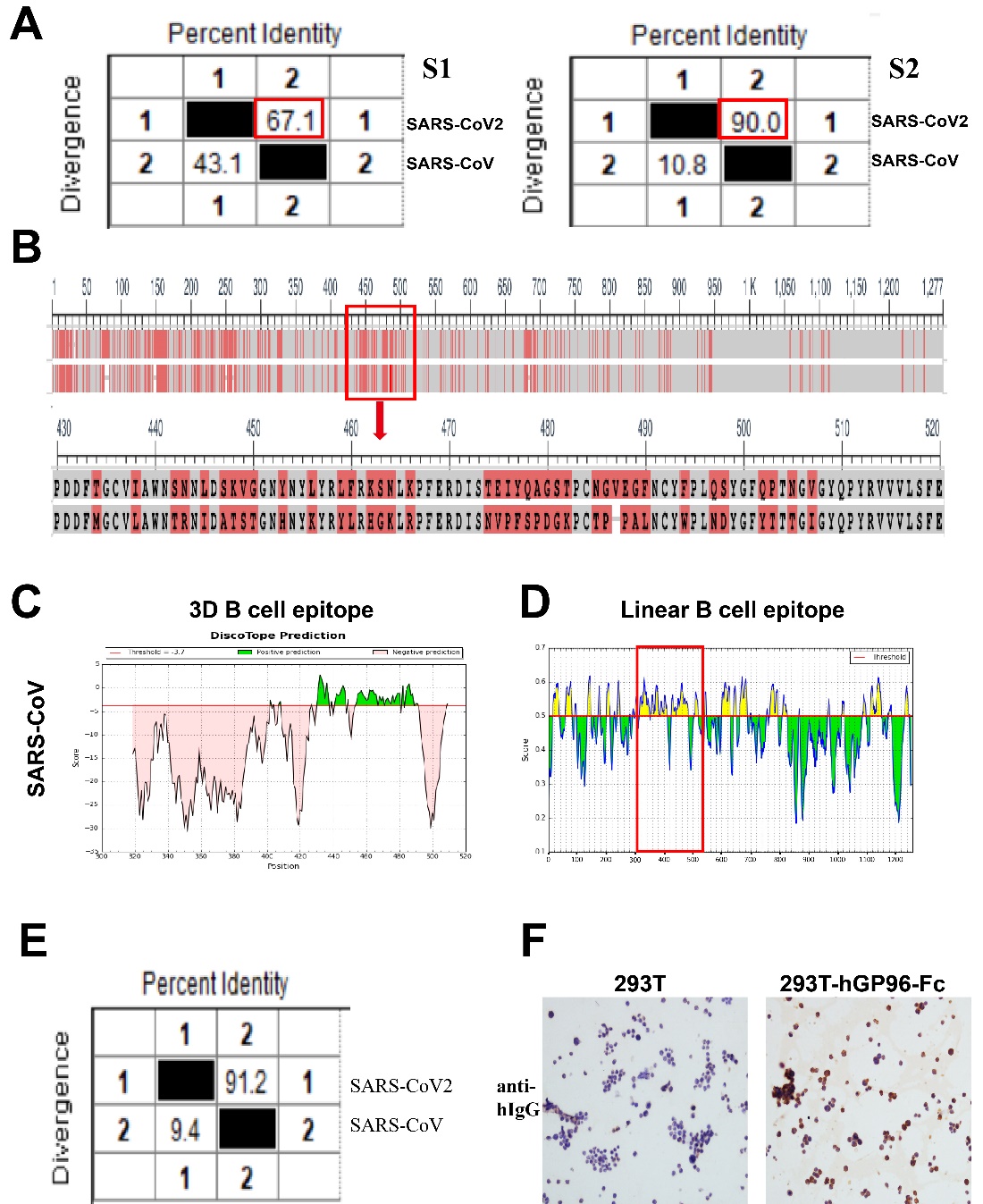

(A) Homology analysis of spike protein (S1 and S2 domain) between SARS-CoV2 and SARS-CoV using MegAlign. (B) Distribution of amino acid variation in two viruses. (C) Potential B cell antigen of RBD domain from SARS-CoV is predicted by Discotope software basing on their 3D structure. (D) Potential linear B cell epitopes of SARS-CoV full S protein are analysed with IEDB database. (E) Homology analysis of nucleocapsid (N) between SARS-CoV2 and SARS-CoV using MegAlign. (E) Protein expression of gp96-hFc fusion protein in stable 293T cells were confirmed with Immunocytochemistry.

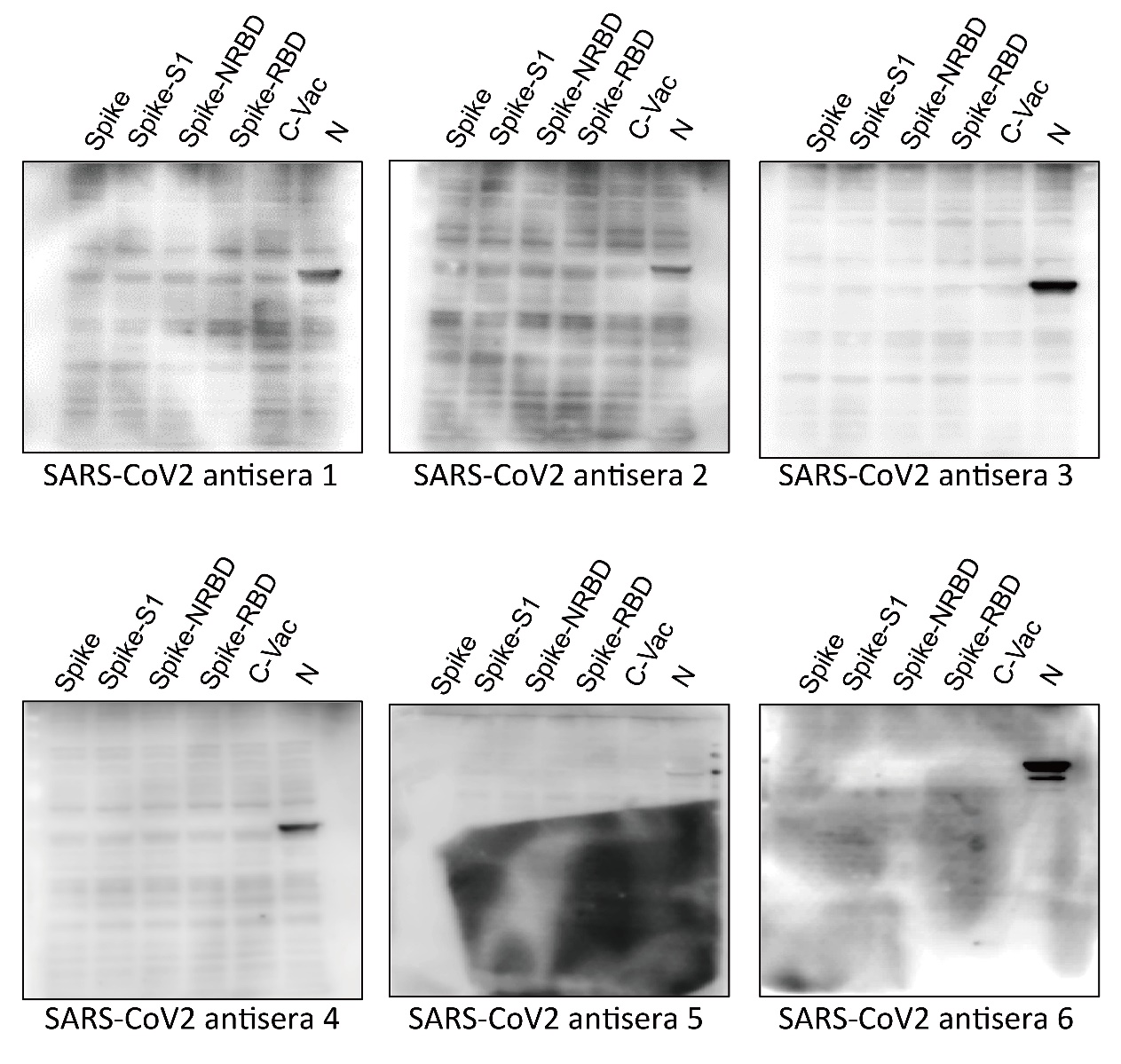
Fig. S2

Fig S2 The pattern of antibodies against the different viral proteins of SARS-COV-2 in the serum of six COVID-19 patients. Antibodies against S, N or truncated N in six SARS-CoV2 antisera were evaluated by Western blot assay. 293T cells were transfected with the plasmids expressing different genes of SARS-COV-2 as indicated, the protein from different groups were probed with the sera of six COVID-19 patients by Western Blot.
